# Phenotypic plasticity promotes balanced polymorphism in periodic environments by a genomic storage effect

**DOI:** 10.1101/038497

**Authors:** Davorka Gulisija, Yuseob Kim, Joshua B. Plotkin

## Abstract

Phenotypic plasticity is known to evolve in perturbed habitats, where it alleviates the deleterious effects of selection. But the effects of plasticity on levels of genetic polymorphism, an important precursor to adaptation in temporally varying environments, are unclear. Here we develop a haploid, two-locus population-genetic model to describe the interplay between a plasticity modifier locus and a target locus subject to periodically varying selection. We find that the interplay between these two loci can produce a “genomic storage effect” that promotes balanced polymorphism over a large range of parameters, in the absence of all other conditions known to maintain genetic variation. The genomic storage effect arises as recombination allows alleles at the two loci to escape more harmful genetic backgrounds and associate in haplotypes that persist until environmental conditions change. Using both Monte Carlo simulations and analytical approximations we quantify the strength of the genomic storage effect across a range of selection pressures, recombination rates, plasticity modifier effect sizes, and environmental periods.

## INTRODUCTION

Balanced polymorphism fosters adaptation in changing environments. As populations continuously adapt from one environment to the other, genetic polymorphism provides a readily available reservoir of adaptive alleles that selection can act upon, and, thus, promotes population persistence (Lande and Shannon 1996; Barrett and Schluter 2008). Despite their role in persistence, evolutionary mechanisms that help maintain genetic polymorphism in temporally changing environments remain poorly understood.

Empirical studies have revealed cases of polymorphism that are subject to temporally varying selection (e.g. Lynch 1987; Cain et al. 1990; Turelli et al. 2001, Bergland et al. 2014). However, the underlying mechanisms maintaining genetic polymorphism in these cases are not either known or confirmed. Theoretical possibilities that predict balanced polymorphism in varying environments include heterozygous advantage (geometric mean overdominance, which cannot occur in haploids; Dempster 1955; Haldane and Jayakar 1963; Gillespie 1973, 1974), overlapping generations with age/stage specific selection or seed banks (Ellner and Hairston Jr 1994; Turelli et al. 2001; Svardal et al. 2011), density regulation with resource competition (Dean 2005; Yi and Dean 2013), or these mechanisms in combination with spatial heterogeneity in selection (Gillespie 1974, 1975; Ewing 1979; Svardal et al. 2015; Gulisija and Kim 2015). Despite these developments, balancing selection due to temporally varying selection has not been widely accepted in population genetic literature probably because early models were criticized due to their failure to maintain polymorphism under genetic drift (Hedrick 1976).

The storage effect, initially recognized in studies of species coexistence in community ecology (Chesson and Warner 1981; Chesson 1985; Chesson 2000), presents a key mechanism underlying balanced genetic polymorphism in populations with overlapping generations and stage-specific selection, or in combination with spatial heterogeneity (Ellner and Hairston Jr 1994; Turelli et al. 2001; Svardal et al. 2015; Gulisija and Kim 2015). The basic idea is that a fraction of the population, in a specific life stage or a patch of habitat where adverse effects of selection are diminished, can “store” polymorphism until conditions change. Although storage effect was originally studied in deterministic framework, Svardal et al. (2011, 2015) demonstrated a storage effect in finite populations experiencing overlapping generations and fluctuating selection, while Gulisija and Kim (2015) demonstrated balanced polymorphisms in finite populations under periodic selection and discrete generations, where a portion of a population is exposed to a decreased magnitude of environmental oscillation. Moreover, Gulisija and Kim (2015) showed that the effect arises even in the presence of geometric mean selective advantage/disadvantage between the competing alleles.

While it is clear that heterogeneous varying selection can lead to the storage effect under various fitness scenarios, past studies have focused on the storage effect arising from age/stage or spatial heterogeneity in varying selection pressures. But another possibility, which we study here, is that a novel mutant allele may experience heterogeneous selection due to its association with different genetic backgrounds that influence the mutant’s phenotypic effect. Thus, diverse genetic backgrounds may take the place of different stages or spatial patches, resulting in a variety of fitness effects on a mutant allele. For example, phenotypic plasticity, the ability of a genotype to produce multiple environmentally induced phenotypes, can modulate the fitness effects of an allele. If there is within-species variation in the plastic response, where some individuals are able to adjust fitness effects of an allele in response to changing environments to a different degree than others, then a plastic subpopulation will experience a different pattern of fitness oscillations than a non-plastic subpopulation. Whether or not heterogeneity in cyclic selection arising from this type of a genomic interaction can promote polymorphism in finite populations is unexplored — and it forms the central question in this study.

Phenotypic plasticity is known to evolve in perturbed or adverse habitats, where it may alleviate fitness-reducing effects of selection (West-Eberhard 2003, Price 2006, Lande 2009, Draghi and Whitlock 2012). Not only can it promote the persistence of a population under adverse conditions, phenotypic plasticity may aid its establishment in novel environments, before genuine genetic adaptation can take place (Galambor et al. 2007, Lande 2009, Fierst 2011). In fact, invasive populations that undergo rapid adaptation typically display a higher degree of plastic response in major fitness components than those that do not rapidly adapt (Lee et al. 2003). However, phenotypic plasticity may also incur fitness cost (DeWitt et al. 1998, Schlichting and Piglicci 1998, Ancel 2000, Lande 2009), such as the cost of development and maintenance of structures and systems involved in osmo- or thermo-regulation (Krebs and Feder 1997). If the cost of plasticity exceeds benefit — e.g., once conditions are no longer adverse and plastic response is no longer needed — plasticity is selected against. Hence, depending on the environmental scheduling, phenotypic plasticity may be adaptive or maladaptive.

Plastic responses may occur due to environmentally sensitive loci and/or “plasticity” modifiers, such as transcription factors or epigenetic modifiers modulating the expression at a target coding sequence (as modeled by Feinberg and Irrizarry 2010, Carja and Feldman 2012, and Draghi and Whitlock 2012). Numerous studies support this epistatic model of plasiticity. Namely, genes other than those primarily responsible for expressing a trait can nevertheless contribute to the trait’s phenotypic plasticity (for a review see Scheiner 1993). Furthermore, mapping studies have identified many quantitative trait loci that modulate phenotypic plasticity, in several model organisms (Stratton 1998, Leips and MacKay 2000; Bergland et al. 2008 Tetard-Jones et al. 2011). When a polymorphic plasticity modifier is not closely linked to the target sequence, alleles at the target locus may recombine to genetic backgrounds yielding different fitness effects, which could emulate a storage effect under varying environments. In particular, plasticity can increase a target allele’s fitness in adverse environments (benefit > cost), but decrease its fitness in favorable environments (cost > benefit). (In this sense, the fitness effect of the modifier here may be equivalent to that of the modifier of genetic robustness (de Visser et al. 2003). Below we use the term plasticity modifier that is broadly defined to include the potential effect of robustness modifier.) In the subpopulation carrying a plasticity modifier, the magnitude of marginal fitness oscillations at a target locus would be smaller relative to that in the subpopulation of non-carriers. Moreover, the modifier locus itself would indirectly experience heterogeneous selection due to pairings with alleles of different fitness effects at the targeted locus. However, whether or not genetic variance at either the target or the modifier locus can be maintained sufficiently to produce balanced polymorphism is unexplored.

Here, we demonstrate balanced polymorphism at both a modifier locus and at a target locus, subject to periodic selection, in finite haploid populations. We call this two-locus effect, whereby linkage disequilibrium generates subpopulations across which alleles experience heterogeneous selection, the “genomic storage effect”. To complement Monte Carlo simulations in finite populations, we also present a deterministic local stability analysis for polymorphic equilibria, in the infinite population size limit. Finally, we examine the effects of population size and mutation rate on the levels of polymorphism at the target and modifier loci preserved by the genomic storage effect.

## MODEL AND METHODS

### Population genetic model

We explore patterns of polymorphism at a plasticity modifier locus and at its target locus under periodic selection in a Wright-Fisher population. We study the model using both Monte Carlo simulation and analytical approximations. At the beginning of each simulation both loci are monomorphic, with the non-plastic allele *(m)* at the plasticity modifier locus and the ancestral allele (a) at the target locus. At time *t*= 0 we introduce a single novel mutant allele at a random one of the two loci: either a derived allele (*d*) at the target locus or a plasticity modifier allele (*M*) at the modifier locus. The remaining monomorphic locus receives a mutant allele in one of the subsequent generations with probability *Nμ* in each, where *N* is a haploid population size and *μ* is a mutation rate. Thereafter no further mutation is allowed in the population. In each discrete generation, *t*, the frequencies of the four haplotypes (*ma, Ma, md*, and *Md*) are subject to (1) deterministic effects of haploid selection, (2) deterministic effects of recombination between the two loci, and (3) finite-population effects of genetic drift. Simulation runs are conducted until alleles at both of the loci are no longer segregating, due to fixation or loss, or until maximum allotted time is reached (see below).

#### Selection

We study a haploid population model to examine the effects of plasticity modifier–target locus dynamics on polymorphism levels. We assume a periodic environment, such that the ancestral allele at the target locus is favored at some times, whereas the derived allele is favored at other times. Although we do not model phenotypes directly, we assume that the modifier allele alters the phenotype at the target allele: in response to adverse conditions the modifier allele triggers a phenotype that it is fitter than the one expressed without the modifier, and here the benefit of phenotypic plasticity exceeds its cost (or any correlated reduction in fitness, due to pleiotropy). However, once the environment becomes favorable there is no longer any benefit to being plastic, and the modifier allele caries only a reduction in fitness due to the cost of plasticity. Thus, the plasticity modifier effectively reduces the strength of periodic selection at the target locus, making the fitness of modifier allele carriers more robust to environmental periodicity as they experience weaker temporal oscillations than non-carriers. As noted in the introduction, the dynamics we analyze extend to any type of modifier that conveys environmental buffering or robustness.

The effect of the modifier allele is described by a single parameter, *p*, which quantifies reduction in the magnitude of varying selection pressure at the target locus, denoted by *s_t_*. We assume that strong density regulation maintains a constant population size, *N* (soft selection, Wallace 1975). And so the expected evolutionary dynamics in the population are described by the relative fitnesses of the four haplotypes, *ma, Ma, md*, and *Md*.

The haplotype frequencies at the beginning of generation *t, x*_am,t_,*x*_aM,t_, *x*_dm,t_, and *x*_dM,t_, are first modified in expectation by selection, producing the expected post-selection haplotype frequencies

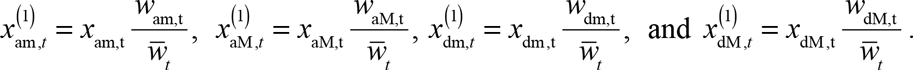

Here

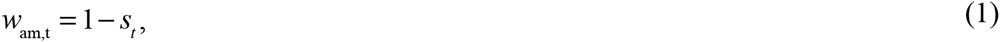

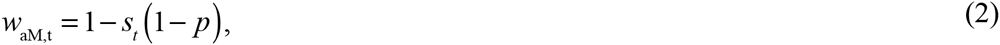

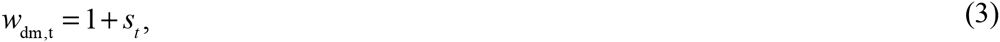

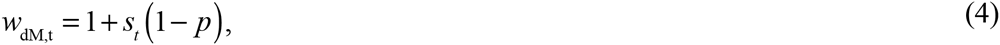

and

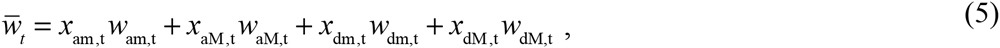

with

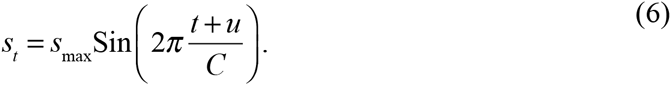

The periodic environment described in Eq. (6) depends upon the parameters *s*_max_, the maximum magnitude of environmental effect; *C*, the length of the oscillating environmental cycle (period); and *u*, which is integer drawn from the uniform distribution between 0 and *C* at time *t* = 0. The random deviate *u* assures that a new mutant allele enters the population at a random phase in the oscillating environment. In the absence of polymorphism at the modifier locus, the derived and ancestral alleles at the target locus have no selective advantage or disadvantage over a cycle of fitness oscillation (that is, 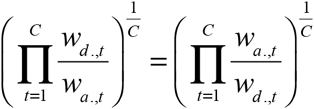 and they are selectively quasineutral in the sense of Hartl and Cook (1973). The modifier locus experiences selection indirectly via its effects on a target allele. In the absence of the polymorphism at the target locus, the plasticity modifier allele is relatively advantageous (that is, 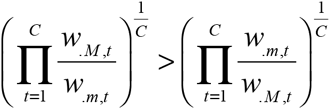). Nonetheless, linkage disequilibrium between the two polymorphic loci might alter fitness effects of alleles. For example, if the modifier allele is associated with a deleterious allele at the target locus, it may experience negative selection.

#### Recombination

Next, the expected haplotype frequencies, 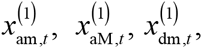 and 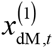, are modified by recombination. Given the coefficient of linkage disequilibrium in generation 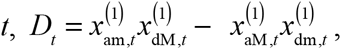, the expected frequencies of four haplotypes following recombination are given by

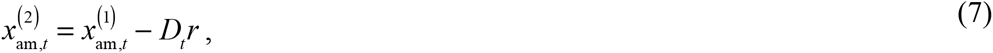

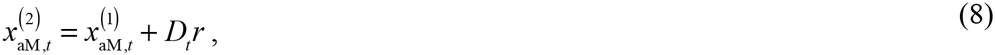

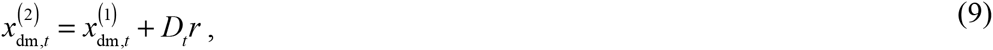

and

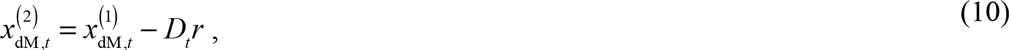

where *r* is the recombination rate between the two loci. The recombination step reduces linkage disequilibrium by a proportion *r*.

**Reproduction and genetic drift:** To describe the effects of genetic drift due to random sampling of gametes in a finite population, we sample *N* individuals from a multinomial distribution using the expected haplotype frequencies after selection and recombination, 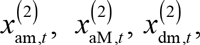 and 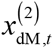 (eqs. (7) – (10)). The resulting sampled frequencies are then used as the starting allele frequencies in the subsequent generation, namely *x*_am,t+1_, *x*_aM,t+1_, *x*_dm,t+1_, and *x*_dM,t+1_.

### Quantifying balanced polymorphism

To quantify the effects of cyclic selection and phenotypic plasticity on the level of genetic polymorphism, we study the expected cumulative heterozygosity at the target locus over time

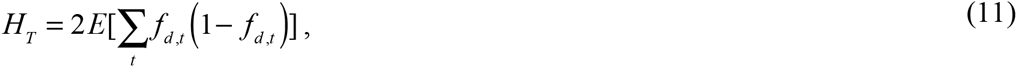

where *f_d,t_ = x_dm,t_ + x_dM,t_* is the frequency of a derived allele at the target locus in the population at time *t*. We also study the expected cumulative diversity at the plasticity modifier locus, *H*_M_, using the frequency of the plasticity modifier locus *f*_M,*t*_ = *x*_aM,*t*_ + *x*_dM,*t*_. From now on, we use the notation *H* for statements pertaining to either *H*_T_ or *H*_M_.

*H* represents expected sum of heterozygosities over a lifetime of a novel mutant (i.e. in the absence of the recurrent mutation), and it provides a simple measure of departure from neutrality that is independent of population size. Under the standard neutral model, *H*_neutral_ = 2 (Kimura 1969). Also, the expected heterozygosity in a haploid population under recurrent mutation equals *NμH* (Kimura 1969) provided the per-site mutation rate, *μ*, and longevity of a new mutant are such that new mutants arise on monomorphic genetic backgrounds (i.e. consecutive mutations do not interfere, *μ* < 1/*NH*). Thus, the ratio *H/H*_neutral_ describes the level of polymorphism relative to that under neutrality, either in the presence or absence of rare mutation.

Here, we use the simulation ensemble-average cumulative diversity, *Ĥ*, as a proxy for polymorphism level under our model. For relatively few simulation runs in which polymorphism persists beyond maximum allotted simulation time, this estimate represents truncated cumulative diversity – that is, *Ĥ ≤ H*.

### Local stability analysis

To complement Monte Carlo simulations in finite populations, we also analyze the population-genetic model above in the limit of infinite *N*, where there are no effects of genetic drift. We set *p* = 1, for convenience, and perform a two-step local stability analysis of the resulting deterministic dynamical system over a range of parameter combinations as described in the Results section. First, we numerically evolve the deterministic difference equations to identify biologically relevant polymorphic equilibrium frequencies, for each parameter set. Next, for each such polymorphic equilibrium we compute the Jacobian matrix of the deterministic system describing allele frequency change over a full cycle of fitness oscillations, and its corresponding eigenvalues, in order to determine whether the equilibrium is locally stable or not.

We numerically determine polymorphic equilibria by evolving the difference equations (1)-(10) starting from numerous boundary conditions where a novel allele (starting frequency = 10^−3^) arises at a locus. The deterministic difference equations are evolved for a burn-in period of 1000 generations and then iterated further, until the frequency of a minor allele at either of the two loci drops below 10^−3^ or until the same sequence of haplotype frequencies (with precision to four decimal points) is repeated in two consecutive cycles of fitness oscillations. Such a repeated sequence of haplotype frequencies represents a two-locus polymorphic equilibrium.

To test the local stability of each identified polymorphic equilibrium, we construct a transition matrix of haplotype frequencies over a full period of fitness oscillations (*C* generations) using the recursion given in equations (1) – (10). The vector of three independent haplotype frequencies in the next generation, X_*t*+1_, given *p* = 1, can be written as

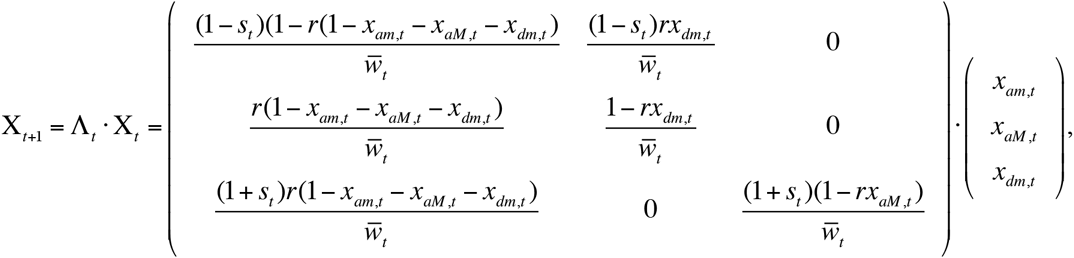

where 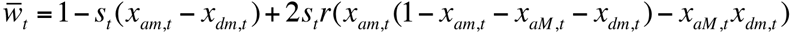 is a mean fitness in the population at the time *t*. If *X_t_* is a vector of equilibrium haplotype frequencies, then it remains unchanged after a full cycle of fitness oscillations

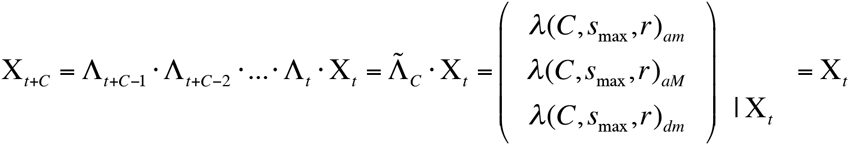

For each such X_*t*_,, we compute the associated Jacobian

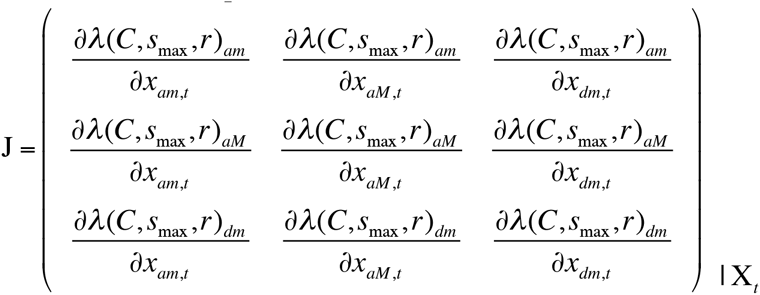

using the central difference formula, 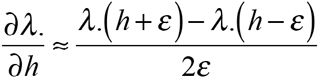 where *h* is a equilibrium haplotype frequency (*x_am,t_, x_aM,t_*, or *x_dm,t_*) at the beginning of the cycle of fitness oscillations. We employ a range of ***ε*** (10^−15^ – 10^−5^) in order to verify the numerical stability of the partial derivatives. Finally, we compute the eigenvalues of *J*. If the absolute value of the leading eigenvalue is less than unity, |*e*| < 1, then the polymorphic equilibrium is locally stable.

### Parameter settings and controls simulations

We study the genomic storage effect across a wide range of fitness effects (*s*_max_ = 0.005, 0.015, 0.05, 0.15, 0.25, or 0.5), oscillating cycle lengths (*C* = 4, 8, 20, 40, or 100), plasticity effects (*p* = 0, 0.25, 0.5, 0.75, or 1), and recombination rates (*r* = 0.0001, 0.01, 0.1, 0.25, or 0.5). All combinations of these parameters were studied via simulations in a haploid populations of constant size *N* = 10^5^, with a single introduction of a mutant allele with probability *Nμ* = 10^−3^ per generation at each locus. For a subset of parameter combinations (as described in the Results section c) we also probed balanced polymorphism in larger populations (*N* = 10^6^) or with recurrent mutation (at rates *Nμ* = 10^−3^ or 10^−1^).

For each parameter set we conducted 5000N independent simulation runs, with a single copy of an allele arising at each locus. We increased the number of simulation runs with *N* to assure that a considerable number of mutants reach high frequency, even in runs with large *N*. However, if both *H_T_* and *H_P_* exceed 20 during the first 10N runs, or if *H_T_* and *H_P_* exceed 100 during the first *N* runs, then we conduct no further simulation runs for that parameter set since, here, a sufficient number of alleles have reached high frequency. Each replicate simulation was run until both loci are no longer polymorphic or otherwise up to 100*N* generations, at which time we record which loci remain polymorphic. The maximum simulation duration of 100N generations was chosen because it is an order of magnitude longer than the maximum duration of polymorphism observed under neutral simulations with genetic drift alone. In the simulations that include recurrent mutation we simulated 1,000 replicate populations of size *N* = 10^5^, with *Nμ* = 10^−1^, and 20,000 replicate populations of size *N* = 10^5^, with *Nμ* = 10^−3^; each of these was run for 100*N* generations.

We also record the time of fixation or loss of all alleles that absorb, in all simulations without recurrent mutation. Along with information about those simulations that continue to maintain diversity at the final time-point, the (100*N*)^th^ generation, these data allow us to quantify the duration of polymorphisms protected by the genomic storage effect.

#### Control Simulations

The cumulative diversity, *H*, may be elevated by mechanisms other than balancing selection. To isolate the effects of heterogeneous cyclic selection from other phenomena that can elevate diversity we conducted a set of control simulations that report polymorphism at each locus in the absence of polymorphism at the other locus. Across a wide range of selection pressures, these control simulations quantify how diversity at the target locus is expected depending on the strength of cyclic selection. Likewise, at the modifier locus these control simulations quantify cumulative diversity that arises from a direct selective benefit of the modifier allele, over the course of its sweep to fixation. In our eventual analysis, we attribute elevated diversity to the genomic storage effect only if cumulative diversity levels at both loci exceed those observed in these control simulations and also exceed the neutral expectation.

#### Control Simulation 1: Target locus in the absence of phenotypic plasticity

In the absence of the modifier allele, cyclic selection results in diversity levels at the target locus, *Ĥ_T_*, that are similar to or less than *Ĥ_neutral_* (Figure 2, bottom left panel), a pattern described in detail in Gulisija and Kim (2015).

#### Control Simulations 2: lasticity modifier locus in the absence of diversity at the target locus

Cumulative diversity levels at the plasticity modifier locus in the absence of polymorphism at the target locus, *Ĥ*_M,control_, typically exceeds *Ĥ*_neutral_ and they range up to 2*Ĥ*_neutral_ (Figure 2, bottom right panel). This elevation in *H* is due to the selective advantage of the modifier allele (see “Population genetic model”), which increases its fixation probability. In these control simulations, the mildly elevated diversity at the modifier locus is not due to balancing selection, but rather to a transient increase in allele frequency as the modifier allele sweeps to fixation, which increases the cumulative measure *Ĥ*_M_. In what follows, we consider only *Ĥ_M_* > *Ĥ*_M,control_ as evidence of elevated diversity caused by balanced polymorphism (as opposed to simply a positively selected sweep at the modifier locus).

## RESULTS

Before describing in detail the results of our simulations and analysis, we briefly summarize the main conclusions they imply. We find that the interplay between a plasticity modifier and a target locus under periodic selection produces a “genomic storage effect” that promotes balanced polymorphism across a wide range of parameter combinations. Persistent polymorphism, characterized by high cumulative diversity (*Ĥ*_T_ > 10*H*_neutral_ and *Ĥ*_M_ > 10*H*_neutral_), occurs exclusively under joint co-variation at the two loci, whereas it never occurs at one locus in the absence of diversity at the other locus. In all parameter regimes that produce such high cumulative diversity we observe long-lived polymorphisms – that is, polymorphisms at both loci that persist for longer than any polymorphism observed in control simulations. The vast majority of these long-lived polymorphisms protected by the genomic storage effect persist for at least 10-times longer than the longest-lived polymorphism observed in the absence of genomic storage (that is, under neutrality, or under either of the control simulations).

The genomic storage effect can have dramatic consequences for the levels of diversity in a population. Consider the ensemble average of heterozygosity at each time point, *h_t_* = 2[*f_t_*(1-*f_t_*)], shown in Figure 1. In the left-hand panel, under cyclic selection but in the absence of the genomic storage effect, the average heterozygosity quickly decreases after introducing the derived allele into the population. The cumulative ensemble average *h_t_*, summed across times (blue lines under the black curve), indicates depressed cumulative diversity at a target locus, *Ĥ*_T_ < 2 (blue colored area), compared to neutrality (yellow, middle figure). On the other hand, in the right hand panel, in the presence of the genomic storage effect, the ensemble average *h_t_* stabilizes at a level far exceeding diversity under neutrality. The resulting elevated value of cumulative diversity, *Ĥ_T_* (indicated in red), indicates a form of balancing selection. Similar patterns of balanced polymorphism are observed across many parameter combinations (see below, and Figure 2).

**Figure 1.**
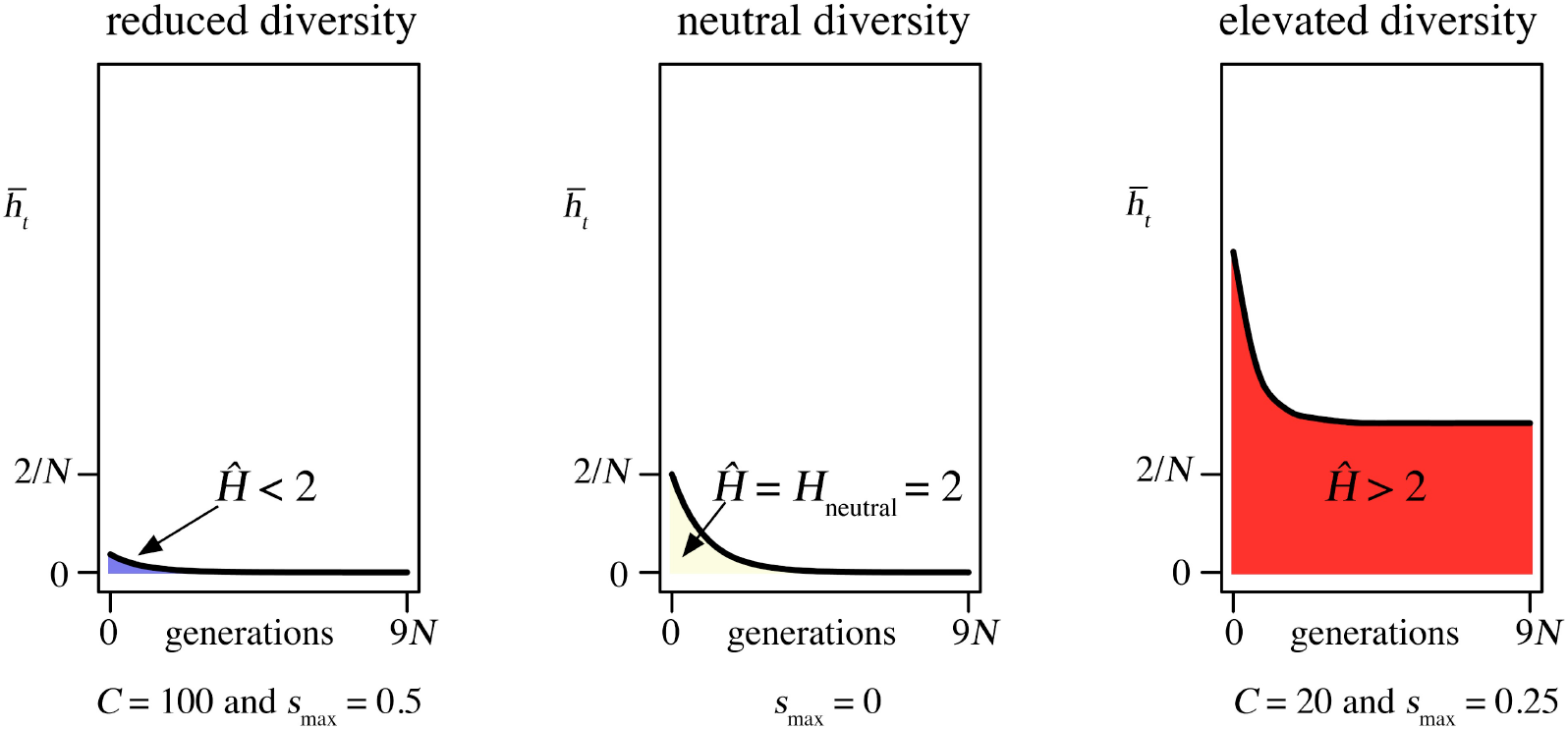
Cumulative heterozygostiy as a measure of diversity. The figure shows the ensemble average heterozygositiy at a target locus under cyclic selection, interacting with an unlinked modifier locus. Cumulative diversity, *H*_T_, is represented in color under the black curves. The heterozygosity was recorded in each generation following the introduction of a single copy of a derived allele. In a parameter set with long and severe seasons (*C* = 100 and *s*_max_ = 0.5), the cumulative diversity is reduced, compared to neutrality. In a parameter set with shorter and milder seasonality (C=20 and *s*_max_ = 0.25) the cumulative diversity is elevated, compared to neutrality. Curves were smoothed via the *loess* fit (span = 0.01 first two panels (5×10^7^ replicates), and span = 0.1 in the last panel (106 replicates)).

**Figure 2.**
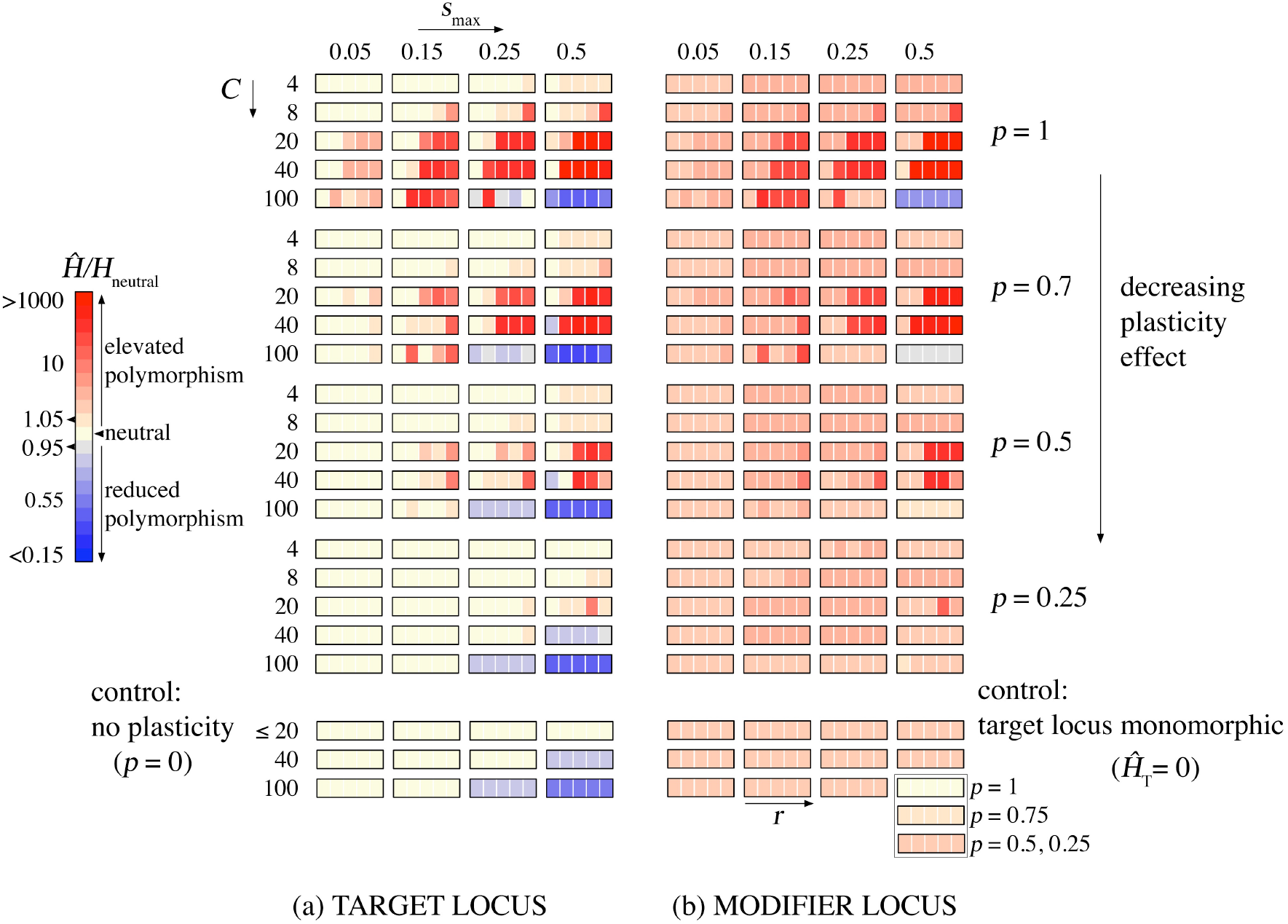
The genomic storage effect promotes polymorphism across a broad range a parameter values. We explore the effects of cyclic selection and phenotypic plasticity on the levels of cummulative diversity at a target locus (*H*_T_ in ‘a’ panel) and at a modifier locus (*H*_M_ in ‘b’ panel) in Monte Carlo simulations of a Wright-Fisher process with selection and recombination. The ensemble-average levels of cumulative diversity are shown as a function of the season length (*C*, increasing vertically within major blocks), the magnitude of periodic seleciton (*s*_max_, increasing horizontally within each panel), the plasticity effect size (*p*, varying vertically across major blocks), and the recombination rate (*r* = 0.0001, 0.01, 0.1, 0.25 and 0.5, varying horizontally within minor blocks). Control simulations, with one or the other locus monomorphic, are down in the bottom blocks of each panel. Only the cumulative divesrity levels that exceed the respective controls represent the genomic storage effect. We simulated an ensemble of 5×10^7^ replicate populations of size *N* = 10^5^, each with a single novel mutant introduction to each of the loci and run for 100*N* generations.

Balanced polymorphism under our model arises in the absence of all known conditions for the maintenance of polymorphism under temporally varying selection. The evolutionary dynamic between a plasticity modifier and a target allele generates an effect analogous to previous models of storage (Chesson and Warner 1981; Chesson 1985; Ellner and Hairston Jr 1994; Turelli et al. 2001; Svardal et al. 2011, 2015; Gulisija and Kim 2015). Modifier and non-modifier genetic backgrounds provide subpopulations in which alleles at the target locus experience different magnitudes/direction of fitness effects, and between which alleles “migrate” by recombination. Likewise, alleles at the modifier locus experience the opposite selection effects as they recombine between different target-locus backgrounds.

In order to definitively confirm the presence of stable polymorphism caused by the genomic storage effect, we complemented Monte Carlo simulations in finite populations with local stability analysis, in the infinite-population size limit. In the following subsections we report in detail (a) finite-population simulation results on the effects of phenotypic plasticity and of cyclic selection on genetic polymorphism, (b) deterministic local stability analysis of two-loci polymorphic equilibria in the infinite-population limit, and (c) the effects of population size and recurrent mutation on the levels of polymorphism in our model.

### (a) Polymorphism under plasticity modifier in periodic environments: simulation results

The genomic storage effect causes balanced polymorphism across a wide range of selection pressures, recombination rates, plasticity modifier effect sizes, and environmental periods (Figure 2). Patterns of balanced polymorphism are similar at the target and modifier loci, as expected, because the maintenance of diversity by genomic storage requires joint polymorphism at both loci. Diversity levels increase with the effect of plasticity (*p*), and with magnitude (*s*_max_) and period (*C*) of fitness oscillations, except under very long seasons with strong selection, in which case alleles fix quickly. With stronger selection and cycle lengths *C* ≥ 8, we can observe simulation runs with both loci still segregating after 100*N* generations, a phenomenon not observed in any simulation runs under control settings, under neutrality, or with any other parameter combination where balanced polymorphism is absent.

Recombination plays a crucial role in the maintenance of allelic variation under the genomic storage effect. Diversity promoted by the genomic storage effect is highest when physical linkage is weak or absent (*r* ≥ 0.25) for short seasons (*C* ≤ 20), or when linkage is moderate (0.0001 < *r* ≤ 0.1) for long seasons (*C* ≥ 40). There is a simple intuition for why the effect of recombination rate on diversity depends on season length. When seasons change quickly, an allele is more likely to persist if it quickly escapes to more favorable background, as under large *r*. On the other hand, when environments tend to be more stable (long seasons), some linkage protects unfavorable alleles from being quickly eliminated from an advantageous genetic background.

The genomic storage effect fails to promote diversity when seasons are long and severe. There are two mechanisms that might disrupt the storage effect in this regime. First, polymorphism is lower with long seasons (*C* = 100) compared to shorter seasons when the allele segregates on a homogenous genetic background (see control simulations in Figure 2). Once a second locus mutates, therefore, the new mutant allele is less likely to arise while polymorphism is present at the initial locus, and cannot therefore benefit from the genomic storage effect. Second, even when joint co-variation occurs, allele frequencies may be pushed to fixation or loss within a single long season, even with the ameliorating effect of the modifier.

The model above uses a sinusoidal fitness function, since we are interested in the effects of phenotypic plasticity on the levels of genetic polymorphism under periodic, predictable environments. Nonetheless, in the Appendix 1 we show that balanced polymorphism arising from the genomic storage effect is robust to stochastic environmental perturbations in season severity and duration, as such perturbations might occur in nature.

### (b) Local stability analysis

The results of local stability analyses, in the infinite-population limit, agree closely with the levels of diversity observed in finite-populations. At each parameter combination where we observed long-lived polymorphism in finite populations, we identified a single, stable two-locus polymorphic equilibrium by local stability analysis (Table S1 in the Appendix 2). Note that such stable polymorphic equilibrium, where selection pushes allele frequencies towards intermediate values, implies a form of negative frequency-dependence. This is analogous to the results of Gulisija and Kim (2015), which demonstrated negative frequency dependence in the context of a spatially heterogeneous cyclic selection.

The range of parameter values for which stable equilibria are observed in deterministic analyses is somewhat larger than the range of balanced polymorphism observed in finite populations. In a deterministic stability analysis, equilibria are identified across the full range of *s*_max_ whereas balanced polymorphism in finite populations is limited to relatively stronger selection. Notably, the leading eigenvalue associated with the Jacobian in our deterministic analysis approaches unity as *s*_max_ decreases (Figure 3, top row). Such leading eigenvalues indicate weak attracting equilibria. In these regimes, selection is not strong enough to maintain polymorphism in the face of the genetic drift in finite population simulations. Moreover, with *C* = 100, even in the cases where attracting stable polymorphic equilibria exist (leading eigenvalue notably < 1), polymorphism perished in finite population since the equilibria included frequencies near zero or one.

**Figure 3.**
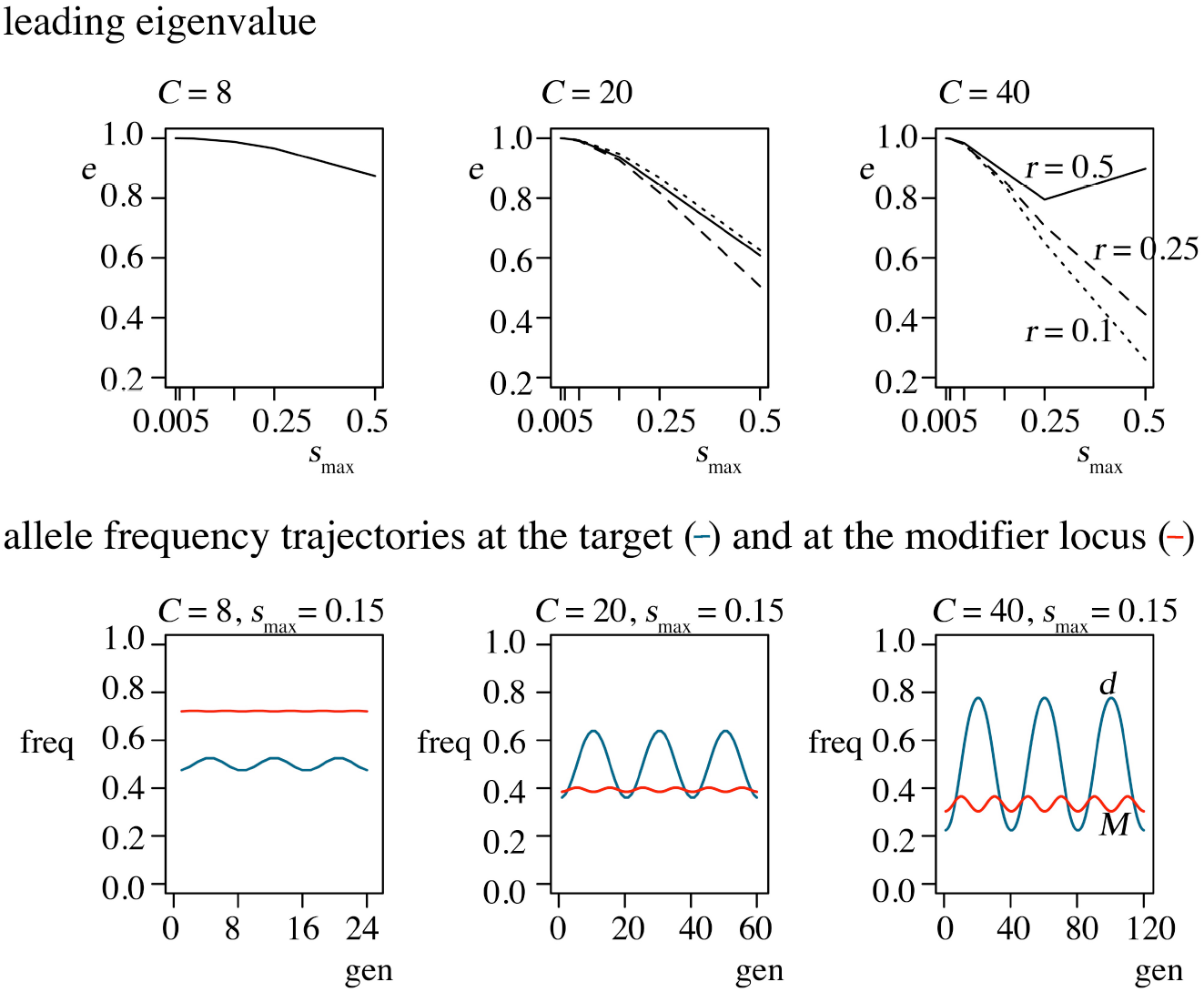
Results of the deterministic local stability analysis. Top panels show the estimated leading eigenvalue of the Jacobian for each equilibrium identified numerically. Bottom panels show sample trajectories of allele frequencies at the target (*d*) and at the modifier (*M*) locus during 3 periods of cyclic selection, at a stable polymorphic equilibrium. *p* = 1 and *r* = 0.5 and in the bottom row.

We briefly describe the form of allele frequency oscillation at stable equilibria (examples are shown in Figure 3, bottom row). The derived allele experiences a single wave and the modifier allele experiences two waves of frequency oscillations. Partly as a result of this, the derived allele tends to oscillate over a wider range of frequencies than the modifier allele. This range is narrower if seasons are short since, in this case, the derived allele experiences shorter periods of selection in the same direction and milder selection (because the majority of individuals carry the modifier allele that buffers environmental effects). At the modifier locus, the non-modifier allele is selected against when paired with a detrimental target allele and selected for when paired with an advantageous allele. The balance between selection effects of different direction at different genetic backgrounds contributes to a relatively narrow allele frequency range at the modifier locus, even when selection is strong over long periods. Long periods of selection against the plasticity modifier allele lead to a reduction in its frequency, while short seasons are characterized by high-frequency modifier alleles.

### (c) The effects of population size and recurrent mutation

While our deterministic analysis in the infinite-population limit predicts balanced polymorphism across the full range of selection coefficients, finite population simulations show no evidence of elevated polymorphism for selection strengths *s*_max_ ≤ 0.015 (data not shown). However, natural populations that experience seasonal succession of generations might be much larger than the one simulated here, *N* = 10^5^ (for example Winkler et al. 2008). This is particularly true since the population size relevant to allele frequencies is not simply the effective population size shaped by demographic history involving bottlenecks, but a typically larger “short-term” effective population size during which adaptation occurs (Karasov et al. 2010). And so we might expect that the genomic storage effect in such popualations promotes diversity over a wider range of selection coefficeints than those discovered in our simulations with *N* = 10^5^. In fact, increasing *N* only 10-fold in our finite population simulations results in balanced polymorphism under far wider range of s_max_ (see example in Figure 4A).

**Figure 4.**
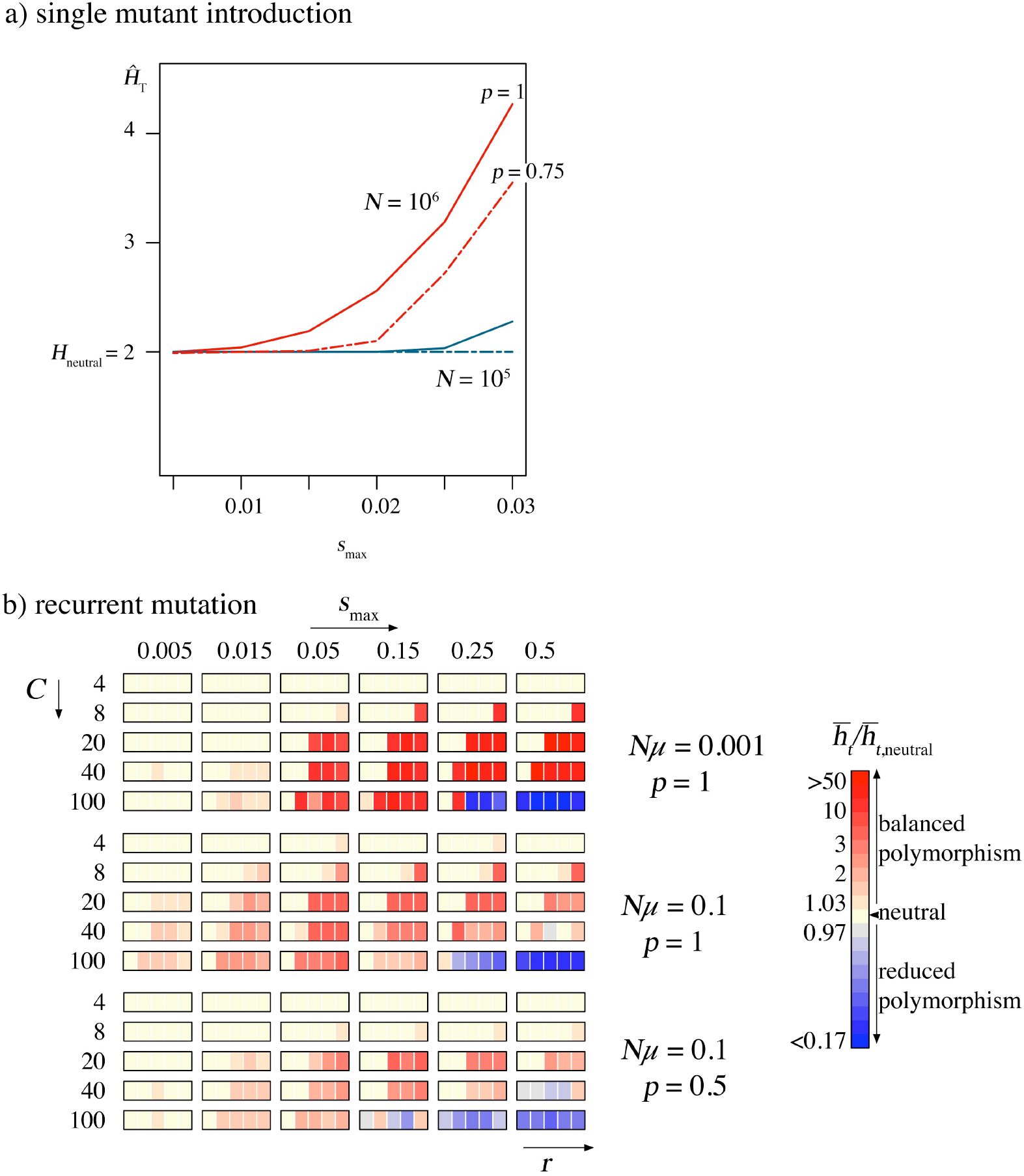
The effect of population size (a) and recurrent mutation (b) on balanced polymorhism under the genomic storage effect. a) Ensemble average heterozygosity at the target locus over time, with *C* = 20, *r* = 0.25, and *p* = 1 (solid line) or *p* = 0.75 (broken line) in a popualtion of size *N* = 10^5^ (blue) or 10^6^ (red), based on the introduction of a single mutant. 10^5^*N* repliacate simulations were each terminated once polymorphism perished or 100*N* generations were reached. b) Ensemble mean heterozygosity, averaged over time, relative to the neutral expection with recurrent mutaiton rate *Nμ* = 10^−3^ and *p* = 1 (top), *Nμ* = 0.1 and *p* = 1 (middle), and *Nμ* = 0.1 and *p* = 0.5 (bottom). We simulated 1,000 replicate populations of size *N* = 10^5^ for *Nμ* = 10^−1^ and 20,000 replicate populations of size *N* = 10^5^ for *Nμ*= 10^−3^, each run for 100*N* generations.

We have seen that polymorphism at one locus (target or modifier) promotes polymorphism at the other locus (modifier or target). And so balanced polymorphism may be more likely if mutation rate is sufficiently large that, during the lifetime of a typical polymorphism, more than one new mutant arises at the other locus. To study this we undertook new simulations with recurrent mutation.

As expected, we found that range of parameters that support balanced polymorphism increases with the recurrent mutation rate, *Nμ*: diversity exceeding the neutral expectation appears across the full range of s_max_ values explored (Figure 4B). However, the extent of elevated heterozygosity relative to neutrality does not increase with *Nμ*, because even neutral heterozygosity quickly saturates to the maximum possible value for large *Nμ*. A similar pattern of balanced polymorphism holds at the modifier locus, although levels of diversity seem lower than under drift in the absence of balanced polymorphism (not shown).

Balanced polymorphism is more likely with large *N* and with recurrent mutation than what we reported in the previous sections. In natural populations, where both of these effects occur, phenotypic plasticity could be a likely precursor to balanced polymorphism and, thus, to rapid adaptation to changing environments.

## INTERPRETATION OF THE TWO-LOCUS DYNAMICS

Here we offer some intuition for the dynamics that produce the sustained polymorphism we have observed under the genomic storage effect. In our model, the target locus experiences heterogeneous periodic selection, due to different genetic backgrounds at the plasticity modifier locus in which target alleles are placed. An allele that is detrimental on a non-modifier background in the current season can recombine (or “escape”) to the modifier background, which buffers its deleterious fitness effect. As the season progresses, selection generates an association between the modifier and the detrimental allele (linkage disequilibrium). Once the season changes, the newly advantageous target allele finds itself linked to the modifier allele, which retards its ability to increase in frequency. However, recombination provides the opportunity for the newly detrimental allele to escape to a less harmful background, and it also allows the advantageous allele to escape to the non-modifier background and thus enjoy full selective advantage.

The plasticity modifier locus is subject to a heterogeneous selection due to the opposite fitness effects its alleles experience when paired with alternative target alleles. Note that, here, heterogeneous selection is different from merely a reduction in selection pressure in a structured population as in the storage effect above: here, selection at the modifier locus has opposite signs on the two different genetic backgrounds. Due to recombination (akin to migration) between the two backgrounds and linkage disequilibrium (akin to subdivision), polymorphism persists until conditions change. In particular, the non-modifier allele recombines to an advantageous target background and experiences increase in frequency. Selection generates an association between the non-modifier allele and the allele that is currently advantageous at the target locus (the coefficient of linkage disequilibrium between them starts from a negative value and then increases to a positive value; see below). Once the season changes, the non-modifier allele finds itself linked to newly detrimental allele, and it is selected against. However, as a non-modifier allele recombines to an advantageous target background (selection leads to reversal in disequilibrium), the non-modifier allele again experiences positive selection. Thus, the non-modifier allele experiences both negative and positive selection within a single season, resulting in two waves of allele frequency changes across an environmental period, as compared to a single wave of frequency change at the target locus.

It is important to understand that the genomic storage effect produces a form of negative frequency-dependent selection. Under the genomic storage effect, the two genetic backgrounds offer both protection and exposure to selection for both alleles at a locus, and alleles survive adverse effects by escaping to a more favorable genetic background. However, the combined effect of both backgrounds does not act equally on both alleles at a locus, but confers a selective advantage to the allele with lower frequency (cf. Gulisija and Kim, 2015). This form of negative frequency-dependence leads to long-lived polymorphism under our model.

## DISCUSSION

This study has revealed a novel mechanism for promoting balanced polymorphism under periodic environmental conditions, which we call the genomic storage effect. The basic idea is that polymorphism under periodic selection can be promoted as recombination allows alleles at two loci to escape more harmful genetic backgrounds and associate in haplotypes that persist until environmental conditions change. This form of genomic storage promotes balanced polymorphism across a wide range of selection pressures, recombination rates, plasticity modifier effect sizes, and environmental periods.

The genomic storage effect we have studied is a new example of a more broad class of storage effects first described in the ecology literature. However, unlike previously recognized storage effects, which arise from age/stage-specific or spatial heterogeneity in selection (Chesson and Warner 1981; Chesson 1985; Ellner and Hairston Jr 1994; Chesson 2000; Turelli et al. 2001; Svardal et al. 2011, 2015; Gulisija and Kim 2015), the genomic storage effect arises as mutation introduces diversity into the two interacting genomic loci, where diversity at one locus serves as heterogeneity generating genetic background to the other. Remarkably, genomic storage promotes diversity in the absence of all of the other known conditions for balanced polymorphism in temporally varying environments, and even in the face of genetic drift in finite population. Below, we address several caveats and implications of this study.

This study proposes the novel idea that linkage disequilibrium driven by selection can promote balanced polymorphism in temporally varying environments. It is important, however, to note that the genomic storage effect we have explored arises from a specific selection regime: (1) a sign change in indirect environmental effect on the non-modifier allele as it recombines between the two genetic backgrounds, i.e. sign epistasis, and (2) a reduction in the magnitude of environmental effect on the target locus when linked to the modifier allele. Balanced polymorphism is recovered only under the two-locus heterogeneous selection. This result points to a novel expectation compared to the single-locus theory, where no balanced polymorphism at either of the loci would be expected when considered on their own. Hence, a single locus theory may be inadequate in describing evolution in periodic environments when several interacting loci are subject to selection. Whereas we have studied the effects of genomic storage in the specific case of two loci, the extent to which single-locus predictions may be altered by other forms of between-loci varying fitness regimes is a topic for future research.

Balanced polymorphism under our model is more likely in large populations and with higher mutation rates, which are both realistic regimes for populations subject to seasonal variation. Nonetheless, the genomic storage effect can also occur under a relatively moderate *Nμ* if the environment imposes stronger fitness effects. The distribution of fitness effects is temporally varying environments is unclear (Eyre-Walker and Keightley 2007). However, it is worth noting that in periodic environments, unlike under directional selection where a new mutant arises in a population close to its optimum, an allele that was fit in one season might become detrimental in the other, giving a strong adaptive advantage to a novel mutant. Moreover, several studies that reported temporally varying polymorphisms implicate strong frequency oscillations (Lynch 1987; Cain et al. 1990; Turelli et al. 2001; Bergland et al. 2014). In particular, Bergland et al. (2014) reported seasonally cycling allele frequencies at hundreds of genomic loci in a sample of flies collected in a single Pennsylvanian orchard. Here, the change in allele frequencies seemed to correspond to (unscaled) selection coefficients ranging from 0.05 to 0.5 per locus per generation. Furthermore, at least some of these cycling polymorphic loci appeared to predate the melanogaster-simulans divergence, suggesting very long-term maintenance of variation. Since strong fitness effects arise under periodic selection, it is plausible that the genomic storage effect could operate in both large and relatively smaller populations in nature.

Gulisija and Kim (2015) argued that a storage effect might arise under a variety of models as long as alleles at a locus are exposed to heterogeneous selection and are close to quasineutral. Indeed, we found that the genomic storage effect is robust to stochastic environmental perturbations in season severity and duration (Appendix 1). While examining the genomic storage effect under different selection models is beyond the scope of this study, we also verified that the genomic storage effect persists across a range of relative benefit and maladaptive effects of plasticity modifier allele, provided there is some part of the season in which the plasticity modifier is beneficial and even in the absence of the maladaptive effect but to much lesser extent (data not shown).

This study not only demonstrates maintenance of polymorphism in periodic environments under a novel mode of balancing selection, but it also makes predictions about the evolution of plasticity. First, our model explains how polymorphism for plasticity may be maintained in periodic environments, without assuming a very large number of underlying loci, as in quantitative-genetic explanations for plasticity. Second, our model implies that, if a population moves to a region with a fixed environment, the non-modifier allele can quickly increase in frequency by selection on standing genetic variation. And so a population could quickly adapt at the target locus, whilst being freed of any cost of plasticity. Thus, our model is in accordance with rapidly adapting invasives originating from perturbed habitats (Ricciardi and MacIsaac 2000; Lee and Gelembiuk 2008).

Finally, our results on genomic storage may have consequences for the evolution of recombination rates in periodic environments. Recombination between the modifier locus and target locus plays a critical role in maintenance of polymorphism under the genomic storage effect. Genomic storage effect, on the other hand, generates cycling linkage disequilibrium. Whether this phenomenon can serve as a basis for selection on the recombination rate remains a topic for future research.

## Acknowledgments

J.B.P. acknowledges support from the David and Lucile Packard Foundation, the U.S. Department of the Interior (D12AP00025), and the U.S. Army Research Office (W911NF-12-1-0552). Y.K. acknowledges support from the National Research Foundation of Korea (2015R1A4A1041997). This research was performed using resources and the computing assistance of the UW-Madison Center for High Throughput Computing (CHTC) in the Department of Computer Sciences. The CHTC is supported by UW-Madison and the Wisconsin Alumni Research Foundation, and is an active member of the Open Science Grid, which is supported by the National Science Foundation and the U.S. Department of Energy’s Office of Science.

